# ALIGaToR: A tool for the automated annotation of immunoglobulin and T cell receptor genomic loci

**DOI:** 10.1101/2025.05.02.651960

**Authors:** Chaim A. Schramm, Simone Olubo, Daniel C. Douek

**Author notes:** Correspondence to: Daniel C. Douek. These authors contributed equally to this work (Chaim A. Schramm, Simone Olubo).

## Abstract

Advances in sequencing technology have made it possible to capture complex immunogenetic loci at a scale that exceeds the capacity for manual annotation. Here we present the Annotator of Loci for ImmunoGlobulins and T cell Receptors (ALIGaToR), an automated pipeline to transfer genetic annotations from a known reference to a novel genomic assembly. We show that ALIGaToR accurately reproduces manually curated annotations and is capable of transferring labels even between distantly related species. Code and documentation for ALIGaToR, including a script reproducing all analyses in this paper, are available at https://github.com/scharch/aligator.

## Introduction

The adaptive immune system is a key defense mechanism used by vertebrates to protect against evolutionarily novel pathogens. In jawed vertebrates, the adaptive immune system is primarily comprises T cells, expressing receptors encoded by the T cell receptor (TR) loci, and B cells, expressing receptors encoded by the immunoglobulin (IG) loci [1]. These loci are characterized by a multiplicity of short, highly similar variable (V), diversity (D), and joining (J) genes, which must be stochastically recombined during lymphocyte development to produce a functional receptor [1]. This complexity makes the IG and TR loci challenging to assemble and annotate [2–8]. The task is nonetheless urgent, as the presence or absence of specific gene polymorphisms have been linked to immune outcomes and disease vulnerability [9–13].

Until recently, immunogenetic annotation of genomic assemblies was done primarily via manual curation [2,14–17]. However, new sequencing technologies [3,18,19] have made it possible to obtain high quality assemblies of immunogenetic loci on a population scale [4,5,20,21] and across a wide variety of species [7,22]. Extracting full immunobiological insight from these data therefore requires an automated tool capable of producing accurate annotations at a similarly high throughput. In addition to V, D, and J genes, the identification of C genes is complicated by substantial structural variability, significant sequence diversity, and, in the IGH, TRA, and TRB loci, the inclusion of kilobase-scale introns between exons [4,8,23,24]. Several existing tools generate such annotations using a variety of search techniques [25–27], but none use standard genomic file formats for output, and only Digger includes coordinates for Recombination Signal Sequences (RSS) and V gene leader exons in its output. More importantly, none of the available tools currently include annotations of C genes.

Here we introduce the Annotator of Loci for ImmunoGlobulins and T cell Receptors (ALIGaToR). ALIGaToR uses entire gene sequences to search and annotate new genomes, thus facilitating the inclusion of C genes and the simple, accurate capture of short D genes [28]. We additionally demonstrate high concordance between ALIGaToR and manually curated gold standard annotations of V and J genes. Annotations are reported in a fully compliant GFF3-formatted file for easy downstream integration with standard genomics tools. ALIGaToR will thus serve as a valuable tool for the high-throughput annotation of newly available immunogenomic data.

## Results

### Pipeline architecture

Currently, the simplest way to identify candidate IG and TR genes remains searching for sequences within a genome that are highly similar to known IG or TR genes. The most widely used repository for fully annotated IG and TR genomic loci is the International ImMunoGeneTics (IMGT) LIGM-DB [29], which uses a flat-file format to store annotations. ALIGaToR includes an ‘extract’ utility that parses the IMGT annotations and produces a corresponding BED file for use in annotating a novel genomic contig (Fig 1).

**Figure 1.**
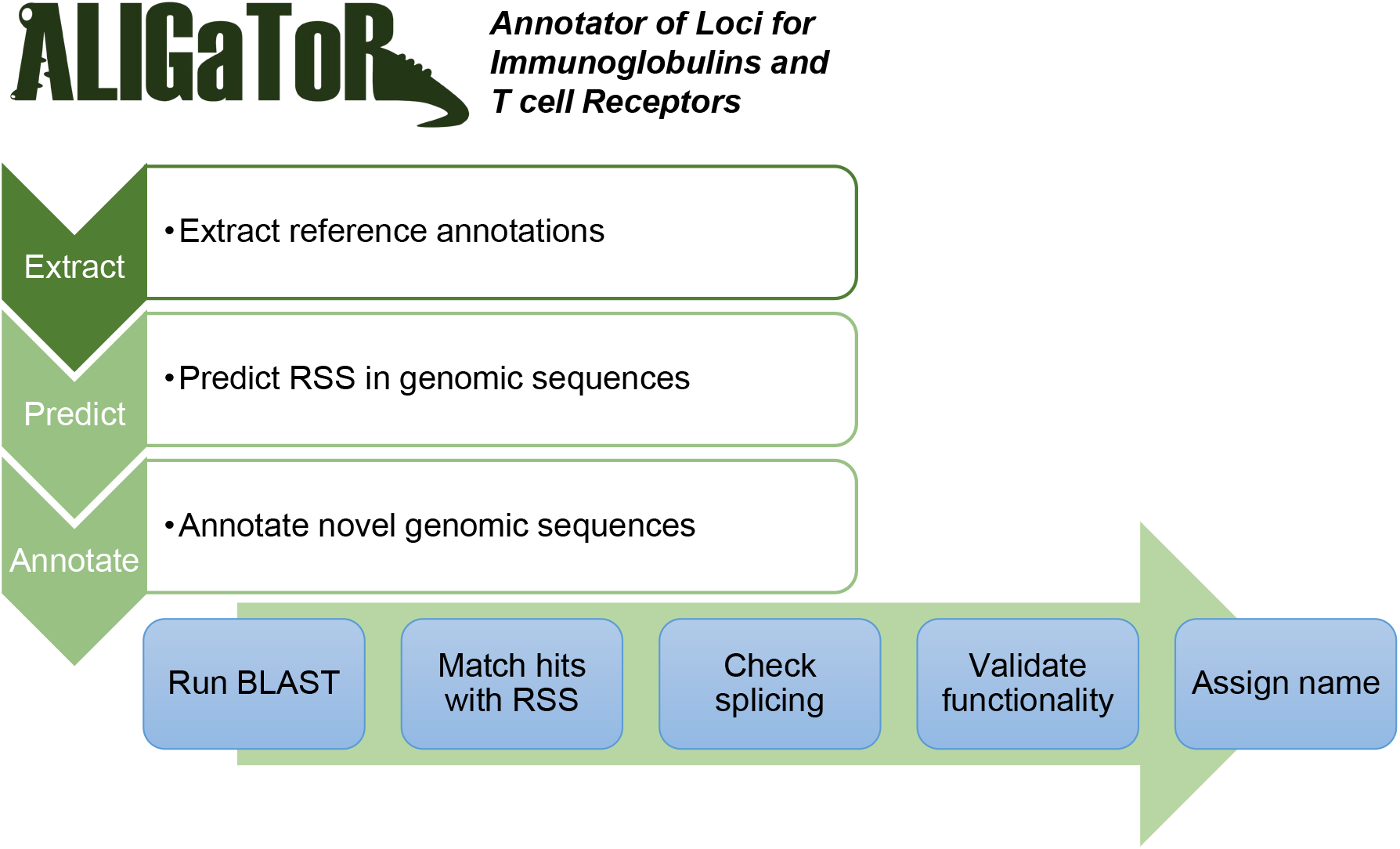
Schematic overview of the ALIGaToR pipeline.

For V, D, and J IG or TR gene to be functional, it must have an RSS for the binding of the RAG enzymes during V(D)J rearrangement [30]. Although some elements of the RSS are conserved, they overall have high sequence variability [31,32], which makes them difficult to detect with tools such as BLAST. ALIGaToR therefore incorporates DNAGrab [33], a tool that uses Bayesian network models to rapidly scan genomic sequences and identify candidate RSS based on the “recombination information content” (RIC) [34,35]. The ‘predict’ command produces a BED file with all of the candidate RSS that can later be matched with V, D, and J genes found with BLAST (Fig 1).

The main module of ALIGaToR is the ‘predict’ function, which sequentially searches for V, D (for IGH and TRB loci), J, and C genes from the reference annotation in the input contig(s). These candidate genes are then matched with predicted RSS positions. Splice sites are verified for V and C genes, and then the coding sequences are checked for stop codons, the presence of a proper start codon (V genes only), and the presence of both conserved cysteine residues in V genes or the conserved tryptophan or phenylalanine residue in J genes. Finally, an identifier is assigned to novel alleles using the first four characters of the SHA-1 hash of the sequence (Fig 1). Genes that are missing expected RSS, canonical GU/AG splicing motifs, or invariant residues are reported by ALIGaToR as “ORF”s, so long as they are complete, contain no internal stop codons, and, for V genes, have a correct start codon.

### IG loci annotation

To validate the ability of ALIGaToR to discover IG genes, we re-annotated genomic assemblies from an Indian-origin rhesus macaque that had previously been manually annotated [16]. For IGH, we used the MF989451 contig and extracted the Mmul_10 reference annotations from IMGT under accession code BK063715 as the starting point for ALIGaToR. We also annotated MF989451 using Digger [27], an alternative tool for automated immunogenetic annotation. While Digger identified a large number of ORF and pseudogenes that were not found by either ALIGaToR or the original curators, the functional genes largely overlapped across the three lists (Fig 2A). Two V genes that were manually identified as functional are instead labeled as ORFs by ALIGaToR, one due to a predicted non-functional RSS (confirmed by Digger), and the other due to a non-canonical GC splice donor after the L-PART1 exon (not checked by Digger) (Fig 2B). ALIGaToR also does not report two IgG constant genes that were manually curated, in both cases because the genes are incomplete due to assembly gaps within the coding sequences. By contrast 4 D gene alleles that were not present in the curated annotations are reported by both ALIGaToR and Digger (Fig 2B).

**Figure 2.**
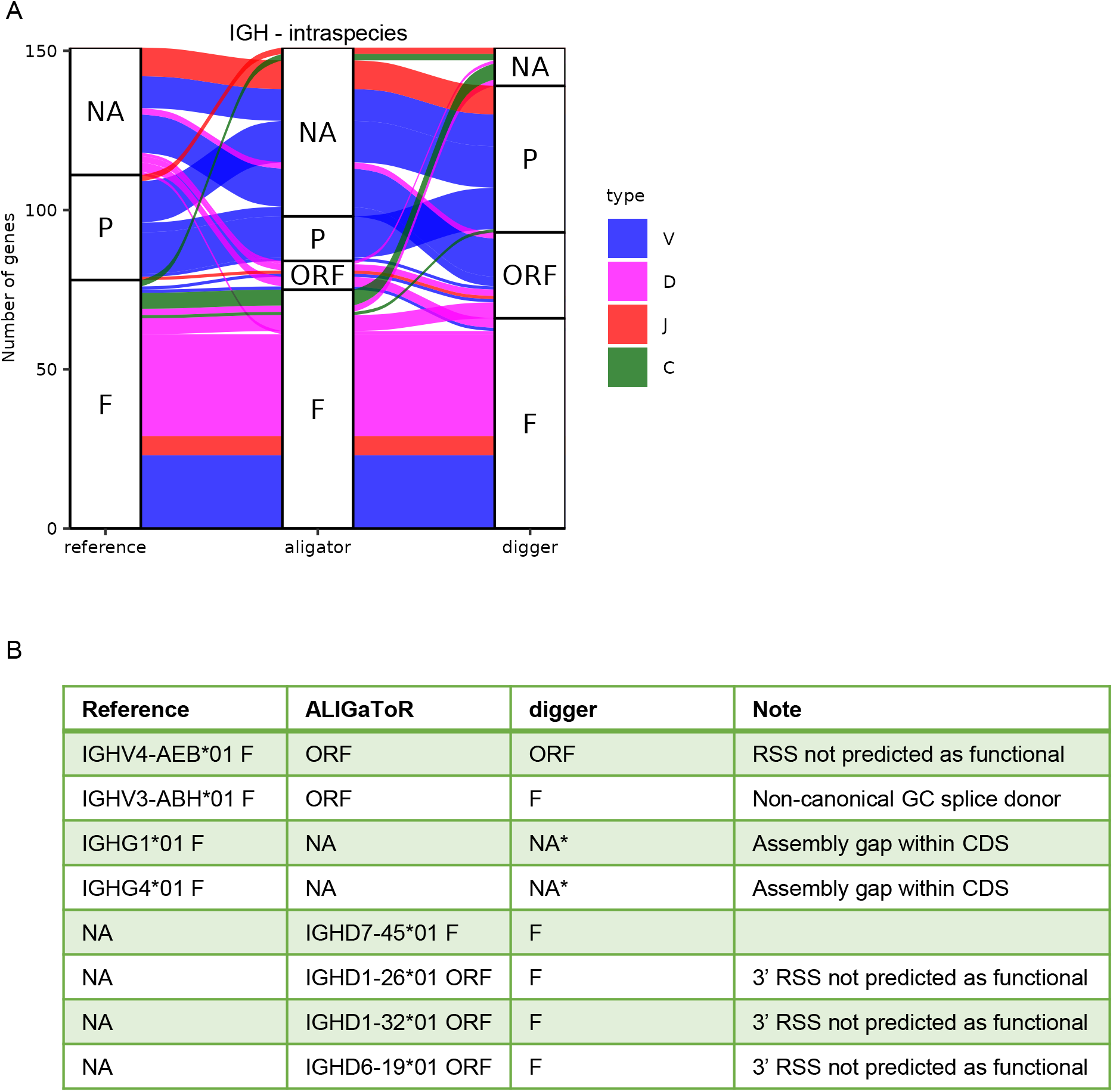
Comparison between unsupervised ALIGaToR annotations of rhesus IGH locus MF989451 with manual curation and automated annotation by digger. (A) Alluvial plot showing the correspondences between levels of functionality assigned by each method. F, functional; ORF, open reading frame; P, pseudogene; NA, not found. Flow colors indicate the gene type. (B) Specific genes that were annotated as functional by one method and where ALIGaToR differed from the manual reference. Note that digger does not currently attempt to annotate constant genes.

For IGK, contig MF989473 was re-annotated using the IMGT Mmul_10 annotations from BK063716. Overall, ALIGaToR closely reproduced the curated annotations (Fig 3A), missing only one V gene with a unique 4 amino acid insertion in CDR1 that may reflect an assembly error in that region of the contig (Fig 3B). Similarly, ALIGaToR finds a duplicated IGKC gene just 600 nt downstream of the primary one, which is almost certainly due to a misassembly that went unnoted by the original authors (Fig 3B). IGL contig MF989453 was re-annotated using the IMGT reference for BK063717, with a similar level of concurrence (Fig 4A). One V gene was flagged as a pseudogene by ALIGaToR due to a frame-shifting deletion in FR3; it is unclear why this gene is annotated in Genbank as having a CDS. By contrast, 5 V genes that do not have CDS annotated in Genbank were labeled as likely functional by ALIGaToR (Fig 4B). ALIGaToR also marked one V gene as an ORF due to a non-canonical splice donor and failed to find one highly divergent V gene (Fig 4B).

**Figure 3.**
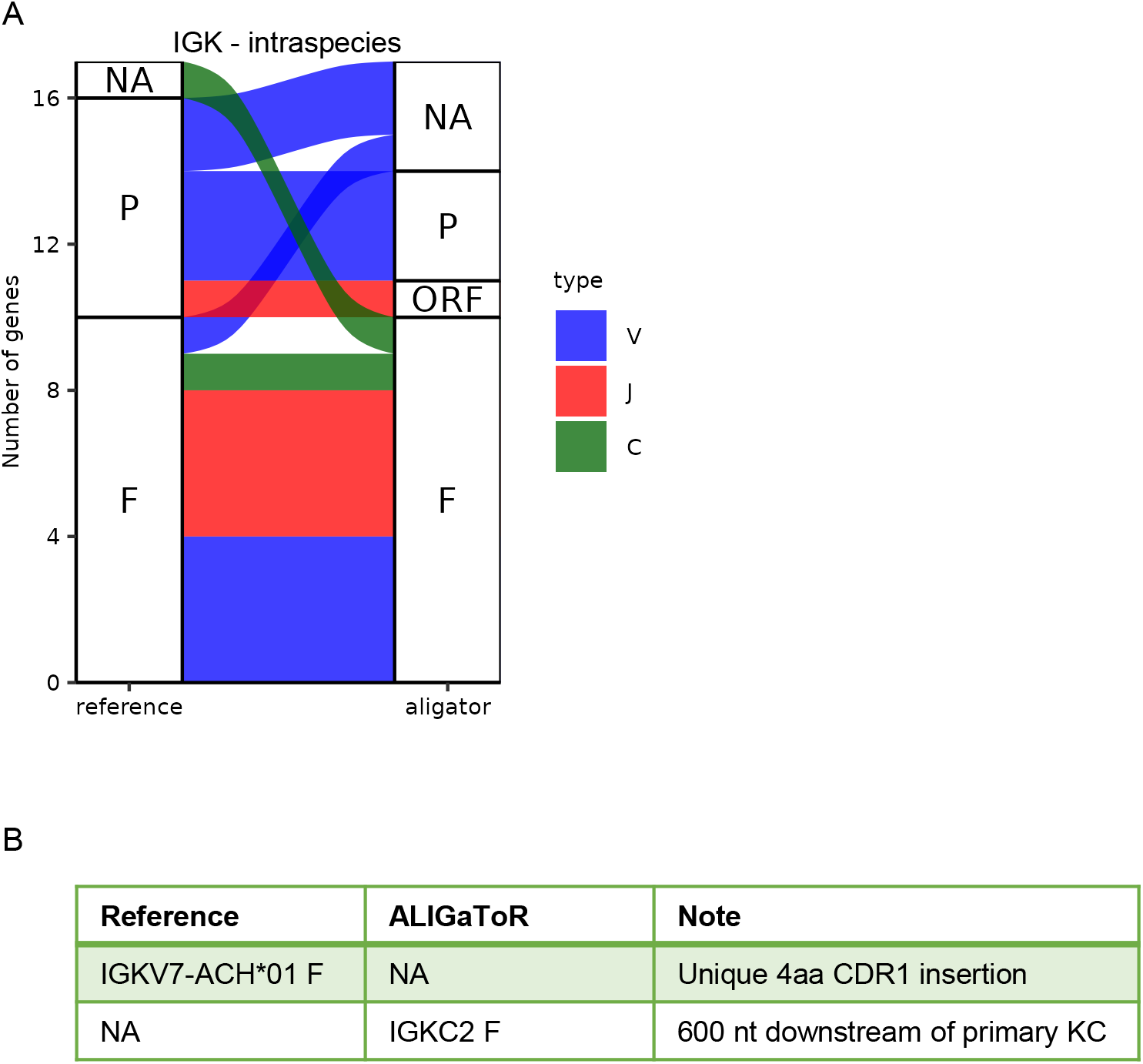
Comparison between unsupervised ALIGaToR annotations of rhesus IGK locus MF989473 with manual curation. (A) Alluvial plot showing the correspondences between levels of functionality assigned by each method. F, functional; ORF, open reading frame; P, pseudogene; NA, not found. Flow colors indicate the gene type. (B) Specific genes were annotated as functional by only one method.

**Figure 4.**
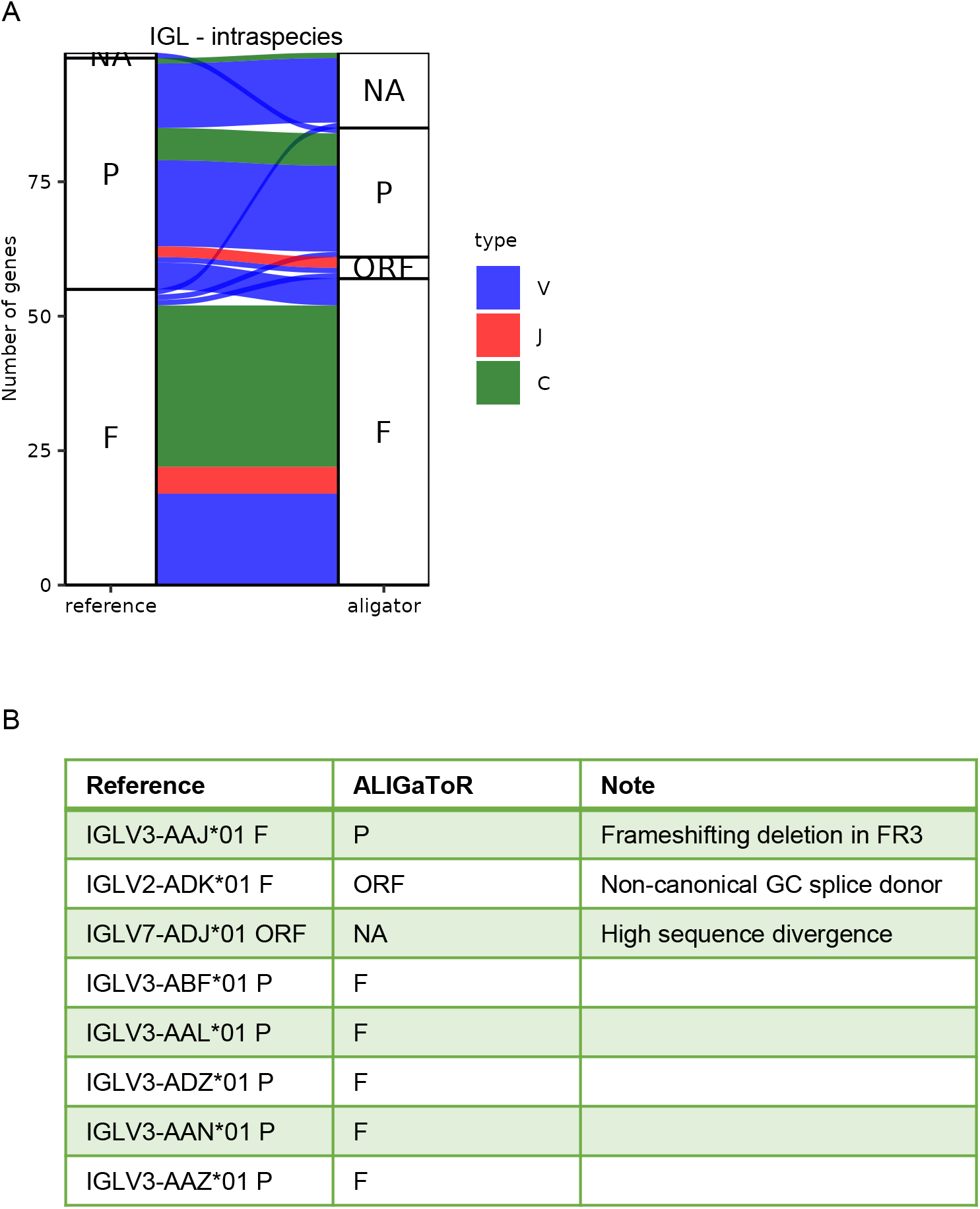
Comparison between unsupervised ALIGaToR annotations of rhesus IGL locus MF989453 with manual curation. (A) Alluvial plot showing the correspondences between levels of functionality assigned by each method. F, functional; ORF, open reading frame; P, pseudogene; NA, not found. Flow colors indicate the gene type. (B) Specific genes were annotated as functional by only one method.

### TR loci annotation

There are currently only 13 species with at least one complete IG or TR locus annotated by IMGT, and only 28 with any V, D, J, or C alleles in the IMGT GENE-DB reference directory, although there are many additional recent annotations in the literature [21,22,36,37]. A common usage case for IG and TR annotation will thus be transfer of orthologous labels from a related species. To test this, we attempted to recapitulate the recent annotation of the ferret TRA/D (BK068537) and TRB (BK068295) loci [38] using the dog genome annotations (BK065026 and BK065025, respectively) as the starting point. These two species are both caniform carnivores, sharing a common ancestor some 40 million years ago [39]. By comparison, humans and macaques share a common ancestor 25-30 million years ago [40]; using the dog TR annotations to re-annotate the ferret TR loci thus provides a reasonably stringent test of the ability to carry gene labels across species. As no RIC model is available for either species, we arbitrarily used the mouse model to predict candidate RSS.

For TRA/D, ALIGaToR recovered 39/61 (64%) functional or ORF TRAV genes, 45/55 (82%) TRAJ genes, and all TRDV, TRDJ, and TRDC genes. ALIGaToR failed to find any TRDD genes or the TRAC gene due to high sequence divergence between the two species (Fig 5A). To confirm that ALIGaToR can detect these genes when starting from a more closely related reference, we also used the same ferret reference annotations as the starting point (Fig 5B). In this case, we recovered all expected genes, though one J gene was marked as pseudogene by ALIGaToR due to a stop codon which can possibly be excised during V-J rearrangement (Fig 5C). Similar results were obtained for TRB, with 13/22 (59%) functional or ORF TRBV genes, 5/11 (45%) TRBJ genes, and 1/2 (50%) TRBC genes, but neither TRBD gene, recovered using the dog annotations as the starting point (Fig 6A). Similar to TRAC1, TRBC1 was missed because the initial BLAST hit was interrupted by low sequence identity in an intron. Using the ferret annotations as the starting point, we recovered all expected genes, with one TRBV gene marked by ALIGaToR as a pseudogene due to a stop codon after the second conserved cysteine, which again could possibly be excised during rearrangement (Fig 6B-C).

**Figure 5.**
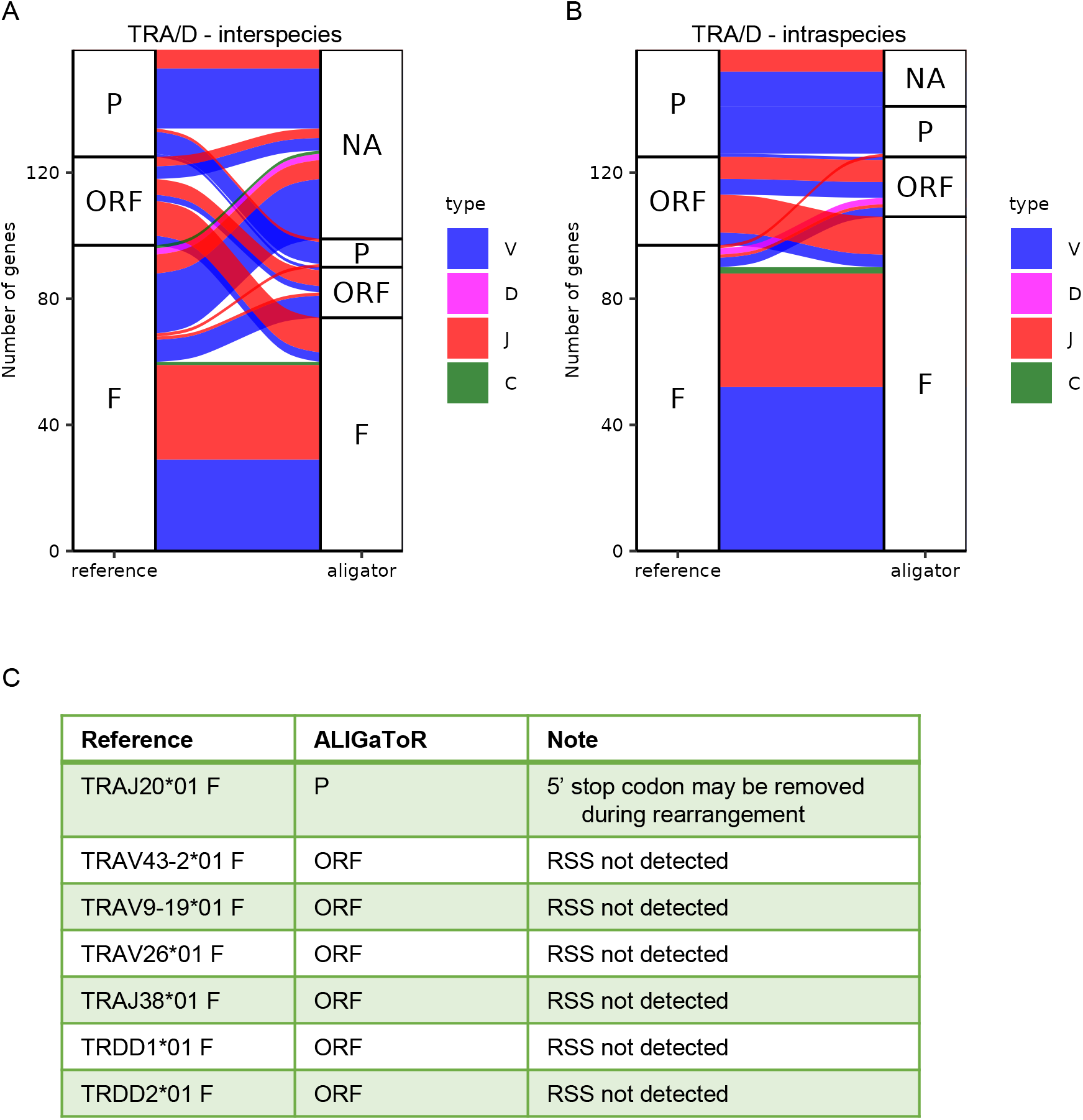
Comparison between unsupervised ALIGaToR annotations of ferret TRA/D locus BK068537 with IMGT curation. (A) Alluvial plot showing the correspondences between levels of functionality assigned by ALIGaToR using the dog TRA/D locus BK065026 as the starting point for annotation. F, functional; ORF, open reading frame; P, pseudogene; NA, not found. Flow colors indicate the gene type. Many functional genes annotated in the reference are not discovered by ALIGaToR due to the high divergence between these two species. (B) Alluvial plot showing the correspondences between levels of functionality assigned by ALIGaToR using the gold-standard IMGT annotations of the same TRA/D locus BK068537 as the starting point for annotation. (C) Specific genes annotated as functional in IMGT that were labeled differently by ALIGaToR.

**Figure 6.**
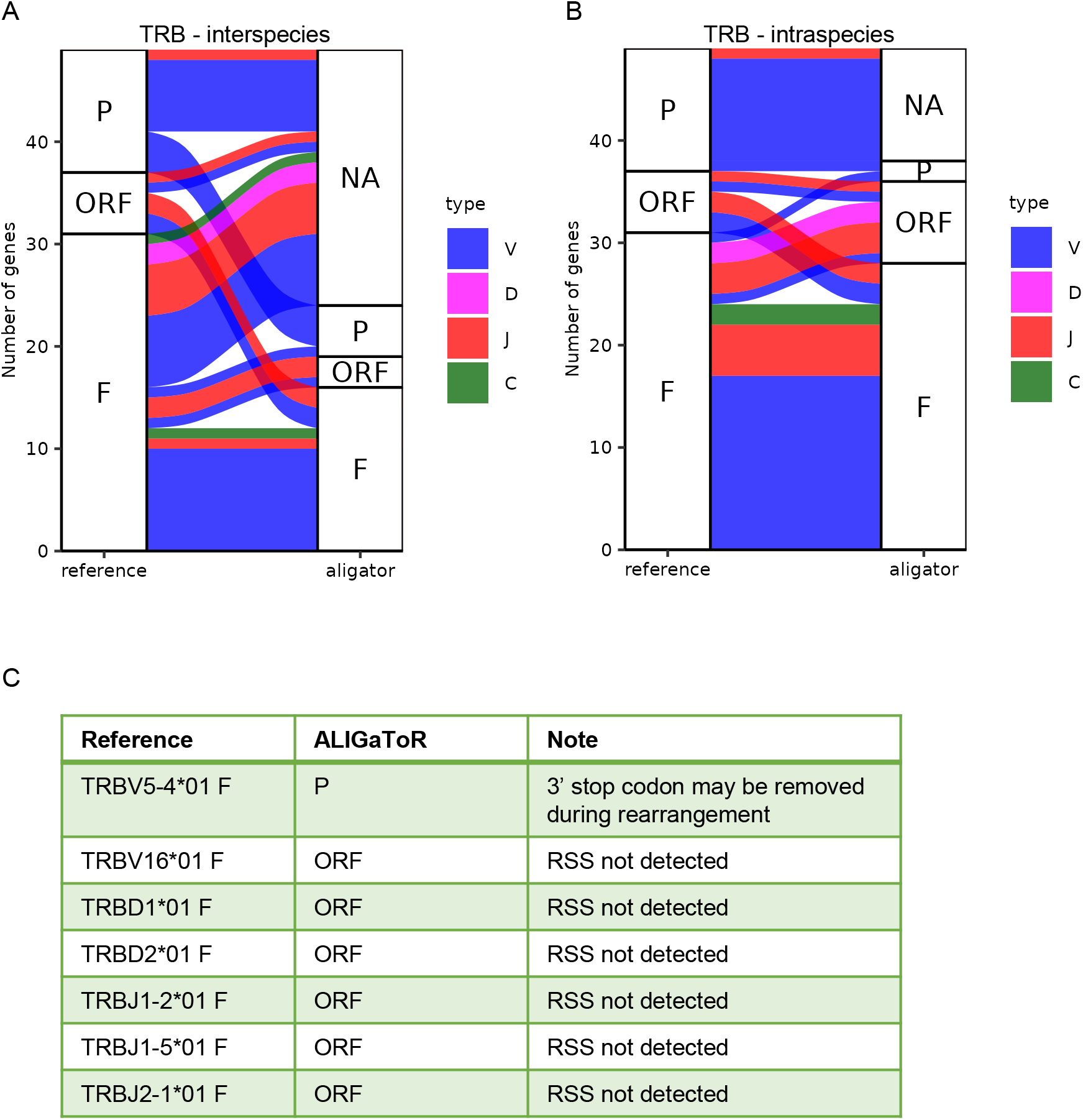
Comparison between unsupervised ALIGaToR annotations of ferret TRB locus BK068295 with IMGT curation. (A) Alluvial plot showing the correspondences between levels of functionality assigned by ALIGaToR using the dog TRB locus BK065025 as the starting point for annotation. F, functional; ORF, open reading frame; P, pseudogene; NA, not found. Flow colors indicate the gene type. Many functional genes annotated in the reference are not discovered by ALIGaToR due to the high divergence between these two species. (B) Alluvial plot showing the correspondences between levels of functionality assigned by ALIGaToR using the gold-standard IMGT annotations of the same TRB locus BK068295 as the starting point for annotation. (C) Specific genes annotated as functional in IMGT that were labeled differently by ALIGaToR.

## Discussion

We have shown that ALIGaToR can faithfully reproduce manually curated immunogenetic annotations given a proper starting point. Even with a fairly distant starting point, ALIGaToR was able to correctly identify the majority of expected genes, with no false positives reported. It may be possible to increase performance on these tasks by raising the maximum e-value for the initial BLAST hits, using an iterative strategy [25,41], and/or starting from multiple reference sources.

Notably, however, the mouse RIC model was unable to detect candidate RSS for several ferret TR genes (Figs 5B-C, 6B-C), despite these being identified by manual curation [38]. This is despite the fact that RIC has previously been found to have a high false-positive rate for reporting RSS [28]. This suggests the need for more specific RIC models or position weight matrices [27], though other tools such as VgenExtractor [42] or Recombination Classifier [43] may also help fill in the gaps. ALIGaToR can use any of these, or others, only requiring that the output be reformatted as a standard BED file.

A unique strength of ALIGaToR is its use of full gene sequences as search input, allowing the accurate capture of C genes. However, this also limits the use of ALIGaToR, as many fewer full genome annotations are available compared to coding sequences, especially for allelic variants of V, D, and J genes. ALIGaToR mitigates the issue by including a functionality to parse gold-standard IMGT annotations, however the lack of fully standardized formatting even within LIGM-DB presents further challenges. Ultimately, ALIGaToR is itself the best solution to this problem, as it generates fully compliant GFF files that can be used as starting points for future annotations. With high accuracy, C gene capture, and interoperability with standard genomics tools, ALIGaToR will be a key tool for high-throughput immunogenetic annotations.

## Methods

Entrez Direct suite [44] was used to obtain reference sequences and annotations from Genbank and a custom perl script (https://github.com/scharch/aligator/blob/master/sample_data/gb2bed.pl) was used to convert the flat-file annotations to BED format. Digger [27] was installed and run following the vignette at https://williamdlees.github.io/digger/_build/html/examples/human_igh.html, with MF989451 replacing IMGT000035 as the target and using the rhesus IMGT reference alleles. The complete ALIGaToR pipeline was run for each reference-target pair. All commands necessary to reproduce the analysis are compiled as a shell script at https://github.com/scharch/aligator/blob/master/sample_data/runTests.sh.

## Data availability statement

All data used here are publicly available under the noted accessions and can be accessed using the commands documented at https://github.com/scharch/aligator/blob/master/sample_data/runTests.sh.

## Study funding

This study was funded in part by the intramural program of the National Institute of Allergy and Infectious Disease, National Institutes of Health.

## Acknowledgements

The authors wish to thank Dr. William S. Gibson for helping to beta-test ALIGaToR and for critical feedback on the manuscript and figures.

## Author Contributions

Conceptualization: CAS

Software: CAS, SO

Supervision: DCD

Visualization: CAS

Writing – original draft: CAS

Writing – review & editing: all authors

## Notes

https://github.com/scharch/aligator

### Competing Interest Statement

The authors have declared no competing interest.

## References

1. Murphy KM, Weaver C. Janeway’s Immunobiology. 9th ed. New Yorkl: Garland Science, 2017.

2. Watson CT, Steinberg KM, Huddleston J et al. Complete Haplotype Sequence of the Human Immunoglobulin Heavy-Chain Variable, Diversity, and Joining Genes and Characterization of Allelic and Copy-Number Variation. The American Journal of Human Genetics 2013;92:530–46.

3. Rodriguez OL, Gibson WS, Parks T et al. A Novel Framework for Characterizing Genomic Haplotype Diversity in the Human Immunoglobulin Heavy Chain Locus. Front Immunol 2020;11, DOI: 10.3389/fimmu.2020.02136.

4. Gibson WS, Rodriguez OL, Shields K et al. Characterization of the immunoglobulin lambda chain locus from diverse populations reveals extensive genetic variation. Genes Immun 2023;24:21–31.

5. Engelbrecht E, Rodriguez OL, Shields K et al. Resolving haplotype variation and complex genetic architecture in the human immunoglobulin kappa chain locus in individuals of diverse ancestry. Genes Immun 2024;25:297–306.

6. Rodriguez OL, Silver CA, Shields K et al. Targeted long-read sequencing facilitates phased diploid assembly and genotyping of the human T cell receptor alpha, delta, and beta loci. Cell Genomics 2022;2:100228.

7. Zhu Y, Watson C, Safonova Y et al. Assessing Assembly Errors in Immunoglobulin Loci: A Comprehensive Evaluation of Long-read Genome Assemblies Across Vertebrates. 2024:2024.07.19.604360.

8. Pospelova M, Voss K, Zamyatin A et al. Comparative analysis of mammalian adaptive immune loci revealed spectacular divergence and common genetic patterns. 2025, DOI:10.1101/2025.04.01.646651.

9. deCamp AC, Corcoran MM, Fulp WJ et al. Human immunoglobulin gene allelic variation impacts germline-targeting vaccine priming. npj Vaccines 2024;9:1–13.

10. Parks T, Mirabel MM, Kado J et al. Association between a common immunoglobulin heavy chain allele and rheumatic heart disease risk in Oceania. Nat Commun 2017;8:14946.

11. Yuan M, Feng Z, Lv H et al. Widespread impact of immunoglobulin V-gene allelic polymorphisms on antibody reactivity. Cell Reports 2023;42:113194.

12. Stephen B, Hajjar J, Sarda S et al. T-cell receptor beta variable gene polymorphism predicts immune-related adverse events during checkpoint blockade immunotherapy. J Immunother Cancer 2023;11:e007236.

13. Safonova Y, Shin SB, Kramer L et al. Revealing How Variations in Antibody Repertoires Correlate with Vaccine Responses., 2021:2021.08.06.454618.

14. Watson CT, Steinberg KM, Graves TA et al. Sequencing of the human IG light chain loci from a hydatidiform mole BAC library reveals locus-specific signatures of genetic diversity. Genes Immun 2015;16:24–34.

15. Ma L, Qin T, Chu D et al. Internal Duplications of DH, JH, and C Region Genes Create an Unusual IgH Gene Locus in Cattle. J Immunol 2016;196:4358–66.

16. Ramesh A, Darko S, Hua A et al. Structure and Diversity of the Rhesus Macaque Immunoglobulin Loci through Multiple De Novo Genome Assemblies. Front Immunol 2017;8, DOI:10.3389/fimmu.2017.01407.

17. Gerritsen B, Pandit A, Zaaraoui-Boutahar F et al. Characterization of the ferret TRB locus guided by V, D, J, and C gene expression analysis. Immunogenetics 2020;72:101–8.

18. Ford M, Haghshenas E, Watson CT et al. Genotyping and Copy Number Analysis of Immunoglobin Heavy Chain Variable Genes Using Long Reads. iScience 2020;23:100883.

19. Zhang J-Y, Roberts H, Flores DSC et al. Using de novo assembly to identify structural variation of eight complex immune system gene regions. PLOS Computational Biology 2021;17:e1009254.

20. Rodriguez OL, Safonova Y, Silver CA et al. Genetic variation in the immunoglobulin heavy chain locus shapes the human antibody repertoire. Nat Commun 2023;14:4419.

21. Peres A, Upadhyay AA, Klein V et al. A Broad Survey and Functional Analysis of Immunoglobulin Loci Variation in Rhesus Macaques. 2025:2025.01.07.631319.

22. Yoo D, Rhie A, Hebbar P et al. Complete sequencing of ape genomes. Nature 2025, DOI:10.1038/s41586-025-08816-3.

23. Ford EE, Tieri D, Rodriguez OL et al. FLAIRR-Seq: A Method for Single-Molecule Resolution of Near Full-Length Antibody H Chain Repertoires. J Immunol 2023;210:1607–19.

24. Jana U, Rodriguez OL, Lees W et al. The human immunoglobulin heavy chain constant gene locus is enriched for large complex structural variants and coding polymorphisms that vary in frequency among human populations. bioRxiv 2025:2025.02.12.634878.

25. Sirupurapu V, Safonova Y, Pevzner PA. Gene prediction in the immunoglobulin loci. Genome Res 2022;32:1152–69.

26. Lin M-J, Lin Y-C, Chen N-C et al. Profiling genes encoding the adaptive immune receptor repertoire with gAIRR Suite. Front Immunol 2022;13, DOI:10.3389/fimmu.2022.922513.

27. Lees WD, Saha S, Yaari G et al. Digger: directed annotation of immunoglobulin and T cell receptor V, D, and J gene sequences and assemblies. Bioinformatics 2024;40:btae144.

28. Safonova Y, Pevzner PA. V(DD)J recombination is an important and evolutionarily conserved mechanism for generating antibodies with unusually long CDR3s. Genome Res 2020;30:1547–58.

29. Giudicelli V, Duroux P, Ginestoux C et al. IMGT/LIGM-DB, the IMGT comprehensive database of immunoglobulin and T cell receptor nucleotide sequences. Nucleic Acids Res 2006;34:D781–784.

30. Gellert M. V(D)J recombination: RAG proteins, repair factors, and regulation. Annu Rev Biochem 2002;71:101–32.

31. Ramsden DA, Baetz K, Wu GE. Conservation of sequence in recombination signal sequence spacers. Nucleic Acids Res 1994;22:1785–96.

32. Hoolehan W, Harris JC, Byrum JN et al. An updated definition of V(D)J recombination signal sequences revealed by high-throughput recombination assays. Nucleic Acids Research 2022;50:11696–711.

33. Merelli I, Guffanti A, Fabbri M et al. RSSsite: a reference database and prediction tool for the identification of cryptic Recombination Signal Sequences in human and murine genomes. Nucleic Acids Res 2010;38:W262–7.

34. Cowell LG, Davila M, Kepler TB et al. Identification and utilization of arbitrary correlations in models of recombination signal sequences. Genome Biology 2002;3:research0072.1.

35. Cowell LG, Davila M, Ramsden D et al. Computational tools for understanding sequence variability in recombination signals. Immunological Reviews 2004;200:57–69.

36. Pursell T, Reers A, Mikelov A et al. Genetically and Functionally Distinct Immunoglobulin Heavy Chain Locus Duplication in Bats. 2024:2024.08.09.606892.

37. Pospelova M, Voss K, Zamyatin A et al. Comparative analysis of mammalian adaptive immune loci revealed spectacular divergence and common genetic patterns. 2025:2025.04.01.646651.

38. Walsh ES, Yang K, Tollison TS et al. Development of ferret immune repertoire reference resources and single-cell-based high-throughput profiling assays. Journal of Virology 2025;0:e00181–25.

39. WesleylHunt GD, and Flynn JJ. Phylogeny of the carnivora: Basal relationships among the carnivoramorphans, and assessment of the position of ‘miacoidea’ relative to carnivora. Journal of Systematic Palaeontology 2005;3:1–28.

40. Stevens NJ, Seiffert ER, O’Connor PM et al. Palaeontological evidence for an Oligocene divergence between Old World monkeys and apes. Nature 2013;497:611–4.

41. Olivieri DN, Gambón-Deza F. Iterative Variable Gene Discovery from Whole Genome Sequencing with a Bootstrapped Multiresolution Algorithm. Comput Math Methods Med 2019;2019:3780245.

42. Olivieri D, Faro J, von Haeften B et al. An automated algorithm for extracting functional immunologic V-genes from genomes in jawed vertebrates. Immunogenetics 2013;65:691–702.

43. Passagem-Santos D, Bonnet M, Sobral D et al. RAG Recombinase as a Selective Pressure for Genome Evolution. Genome Biol Evol 2016;8:3364–76.

44. Kans J. Entrez Direct: E-utilities on the Unix Command Line. Entrez Programming Utilities Help [Internet]. National Center for Biotechnology Information (US), 2025.

